# Genome-wide analysis of the annexin gene family in alfalfa (*Medicago sativa* L.) and functional characterization of *MsANN22* under drought stress

**DOI:** 10.64898/2026.10.21.760862

**Authors:** Zhengjun Sheng, Guangzhi Jiang, Jiayin Wang, Huan He, Pingping Du, Hongfei Li, Zhuang Meng, Qinglong Shao, Hongbin Li, Quanliang Xie

**Author notes:** These authors contributed equally (Zhengjun Sheng, Guangzhi Jiang, Jiayin Wang). Corresponding authors (Q. Xie), (H. Li), (Z. Meng).

## Abstract

Annexins are a conserved family of calcium-dependent phospholipid-binding proteins that participate in diverse plant stress responses. However, the annexin gene family in Medicago sativa and the molecular mechanisms underlying annexin-mediated drought tolerance remain largely unexplored. In this study, 73 *MsANN* genes were identified in the alfalfa genome. Their phylogenetic relationships, synteny, conserved motifs, gene structures, and promoter cis-acting elements were systematically analyzed. Synteny analysis revealed 75 segmental/whole-genome duplication-associated gene pairs but no tandem duplications, indicating that large-scale duplication was the major driver of *MsANN* family expansion. The *MsANN* promoters contained ABA- and drought-responsive elements. Expression profiling identified MsANN22 as a strongly drought-responsive gene encoding a cytoplasm-localized protein. *MsANN22* overexpression enhanced drought tolerance in both Arabidopsis thaliana and alfalfa: in Arabidopsis, *MsANN22* overexpression promoted seed germination under osmotic stress, whereas in alfalfa it enhanced antioxidant capacity, with superoxide dismutase and peroxidase activities reaching 1.38- and 1.70-fold those of wild-type plants, respectively. Consistently, *MsANN22*-overexpressing alfalfa accumulated approximately 20% less malondialdehyde and exhibited reduced H O and O • accumulation, as indicated by DAB and NBT staining. Yeast two-hybrid and luciferase complementation assays further supported physical interactions of MsANN22 with MsIAA and MsPR-10. These findings provide insight into the molecular mechanism by which *MsANN22* mediates drought tolerance and offer a candidate gene for improving drought resistance in alfalfa.

## 1. Introduction

Annexins are a multigene family of Ca²□-dependent phospholipid-binding proteins that are widely distributed across plants and animals (Rescher and Gerke, 2004). They have been identified in numerous plant species (Cantero *et al*., 2006; Zhang *et al*., 2015; He *et al*., 2019; Wei *et al*., 2022). Compared with animal annexins, plant annexins generally possess shorter N-terminal regions and simpler domain architectures, while retaining conserved Ca²□- and phospholipid-binding properties associated with membrane-related functions (Gerke and Moss, 2002). In addition, some plant annexins exhibit peroxidase or ion-channel activity. They participate in physiological processes such as growth, differentiation, and secretion, and contribute to responses to drought, salinity, oxidative stress, and tissue damage (Clark *et al*., 2012; Faizan *et al*., 2024; Xie *et al*., 2025).

*MsANN2* was the first annexin gene identified in alfalfa, and its expression was shown to be induced by drought, salinity, alkaline stress, and exogenous ABA (Kovács *et al*., 1998). Under the genome-wide nomenclature system based on chromosomal positions adopted here, the previously reported *MsANN2* corresponds to *MsANN72*. The two encoded protein sequences share more than 99% identity. Subsequent studies have further demonstrated important functions of annexins in leguminous species. In *Medicago truncatula*, *MtANN2* is highly expressed in roots and specifically expressed in the lateral root primordia of seedlings. A T-DNA insertion mutant of its *Arabidopsis* homolog *AtANN2* exhibits reduced growth and produces fewer lateral roots. *MtANN2* overexpression partially rescues this phenotype and improves seed germination and seedling growth under salt stress, indicating that *MtANN2* participates in root development and salt stress responses (Liu *et al*., 2022). In alfalfa, proteomic analysis identified 52 differentially expressed proteins in roots under salt stress. Ca²□ signaling-related proteins, including calmodulin and annexin *MsANN2* was significantly upregulated. Heterologous expression of *MsANN2* enhanced salt tolerance in *Arabidopsis*, while transgenic alfalfa exhibited increased survival, enhanced antioxidant enzyme activities, and reduced membrane damage under salt stress (Gao *et al*., 2024).

Accumulating evidence indicates that annexin overexpression can enhance plant tolerance to multiple abiotic stresses, including heat, drought, and salinity. For example, *OsANN1* overexpression improves heat tolerance in rice by regulating redox homeostasis (Qiao *et al*., 2015), whereas *GhANN1* overexpression enhances salt tolerance in cotton by modulating ABA metabolism and phenylpropanoid biosynthesis (Zhang *et al*., 2021). In *Arabidopsis*, *AtANN1* and *AtANN4* cooperatively regulate salt and drought tolerance (Huh *et al*., 2010). *AnnSp2* overexpression enhances drought and salt tolerance in tomato (Ijaz *et al*., 2017), whereas *OsANN3* overexpression improves drought tolerance in rice, while suppression of *OsANN3* increases drought sensitivity (Li *et al*., 2019). Constitutive *AnnBj1* expression also enhances tolerance to multiple stresses and improves fiber quality in cotton (Divya *et al*., 2010). Annexin expression is further regulated by phytohormones. In tobacco, *NtAnn12* is specifically expressed in roots following IAA induction (Baucher *et al*., 2011), whereas ABA and GA induce distinct annexin family members in tomato (Lu *et al*., 2012). Ca²□ is an essential second messenger that regulates stress-responsive gene expression (DeFalco *et al*., 2010). As calcium-responsive proteins, annexins participate in Ca²□ signaling-associated physiological processes. Onion annexin promotes tissue growth by modulating calcium and ion homeostasis (Chen *et al*., 2013), whereas cotton annexin p34 inhibits callose synthase activity in a Ca²□-dependent manner (Andrawis *et al*., 1993). These findings suggest that annexins may function as molecular links between Ca²□ signaling and ROS homeostasis.

Nevertheless, how annexins cooperate with other proteins to transmit Ca²□ signals during stress responses remains poorly understood. In particular, the annexin gene family has not been systematically characterized in alfalfa, and its role in drought tolerance remains unclear. Here, we performed genome-wide identification and evolutionary analysis of the alfalfa annexin family, identified the drought-responsive gene *MsANN22* from transcriptome data, and characterized its function through heterologous overexpression in *A. thaliana*, stable transformation in alfalfa, physiological and biochemical assays, and protein–protein interaction screening. Our results provide a basis for dissecting annexin-mediated stress responses in legumes and identify *MsANN22* as a candidate gene for drought-tolerance breeding in alfalfa.

## 2. Materials and Methods

### 2.1 Identification of the *MsANN* Gene Family in Alfalfa

The genome, protein, and coding sequences of the alfalfa cultivar ‘Xinjiang Daye’ were downloaded from the Figshare database (https://figshare.com/). Genome annotation files for *Arabidopsis thaliana* were obtained from the TAIR database (https://www.arabidopsis.org/index.jsp). Amino acid sequences of *Arabidopsis* annexin genes were retrieved and used as queries in BLASTP searches against the alfalfa protein dataset, with an E-value cutoff of ≤ 10□□. In parallel, the hidden Markov model (HMM) profile of the conserved annexin domain (PF00191) was downloaded from the Pfam database (http://pfam.xfam.org), and HMMER was used to search the alfalfa proteome at the same threshold. Both approaches independently identified the same set of 73 candidate sequences. All candidates were further verified using the NCBI Conserved Domain Database (CDD, https://www.ncbi.nlm.nih.gov/cdd) and SMART (http://smart.embl-heidelberg.de/), and all 73 sequences were confirmed to contain the annexin domain (PF00191, E-value≤ 10□□), with no non-annexin proteins retained. The physicochemical properties of the MsANN proteins, including amino acid length, molecular weight, isoelectric point, and grand average of hydropathicity (GRAVY), were calculated using the Protein Parameter Calc module in TBtools-II. Subcellular localization was predicted using WoLF PSORT (https://wolfpsort.hgc.jp/).

### 2.2 Phylogenetic Analysis of *MsANN* Proteins in Alfalfa

Genome data for *Medicago truncatula*, *Oryza sativa*, and *Glycine max* were downloaded from the Ensembl Plants database (https://plants.ensembl.org/) and Phytozome (https://phytozome-next.jgi.doe.gov/). Annexin genes in these species were identified using the same strategy as described for alfalfa, yielding 20, 10, and 26 members, respectively. Multiple sequence alignment was performed using ClustalW with default parameters. A phylogenetic tree was constructed in MEGA X using the maximum likelihood (ML) method with the JTT substitution model and 1,000 bootstrap replicates.

### 2.3 Protein Structure, Conserved Motif, and Chromosomal Localization Analyses

MsANN protein sequences were uploaded to the MEME online server (https://meme-suite.org/meme/) (Bailey *et al*., 2009) to identify conserved motifs, with the maximum number of motifs set to 10. Gene structures, conserved motifs, and the phylogenetic tree were analyzed and visualized using TBtools-II (Chen *et al*., 2020). Chromosomal localization was determined by importing the *MsANN* gene identifiers and GFF3 annotation file into TBtools-II.

### 2.4 Promoter Cis-Acting Element and Synteny Analyses

The 2,000-bp sequences upstream of the translation initiation sites (ATGs) of the *MsANN* genes were extracted as promoter sequences using TBtools-II based on the alfalfa genome and GFF3 annotation files. Cis-acting elements were predicted using the PlantCARE online platform (http://bioinformatics.psb.ugent.be/webtools/plantcare/html/) (Lescot *et al*., 2002). The predicted elements were compiled and visualized in TBtools-II. Intragenomic and interspecific syntenic relationships were analyzed using the One Step MCScanX-Super Fast module (Wang *et al*., 2012) with the corresponding genome and GFF3 annotation files of alfalfa, *M. truncatula*, soybean, rice, and *Arabidopsis*. For duplicated *MsANN* gene pairs, Ka, Ks, and Ka/Ks values were calculated using the Ka/Ks Calculator module in TBtools-II, and the results were visualized using TBtools-II.

### 2.5 Expression Analysis of *MsANN* Genes in Alfalfa

Raw transcriptome data were obtained from the NCBI database (https://www.ncbi.nlm.nih.gov) under BioProjects PRJNA1019078 (drought treatment) and PRJNA907206 (salt and jasmonic acid treatments. The data were converted to FASTQ format using SRA-Toolkit v2.9 (NCBI, USA). Raw reads were quality-controlled and filtered using Trimmomatic v0.39 (Bolger *et al*., 2014). Clean reads were aligned to the alfalfa reference genome using HISAT2 v2.2.1 (Kim *et al*., 2019), and gene expression was quantified using StringTie v2.1.3 (Pertea *et al*., 2015). Expression levels were reported as fragments per kilobase of transcript per million mapped reads (FPKM). Gene expression heatmaps were generated using TBtools-II.

### 2.6 Plant Materials and Treatments

The alfalfa cultivar ‘Xinjiang Daye’ and *A. thaliana* (Col-0) seeds were provided by the Key Laboratory of Xinjiang Phytomedicine Resources and Utilization, Ministry of Education, Shihezi University. Alfalfa seedlings were cultivated until their root systems were fully developed and were then transferred to Hoagland nutrient solution for 7 days of pre-culture. Plants were maintained at 25□ under a 16-h light/8-h dark photoperiod and a light intensity of 6,000 lx. The seedlings were subsequently treated with 50 μM methyl jasmonate (MeJA), 100 μM abscisic acid (ABA), or 15% (w/v) polyethylene glycol 6000 (PEG 6000). MeJA and ABA were applied by foliar spraying until runoff, whereas PEG 6000 was added directly to the Hoagland nutrient solution to simulate osmotic stress. Leaf and root samples were collected at 0, 3, 6, 12, and 24 h after treatment, with three biological replicates per time point. Additionally, leaf, stem, and root tissues were harvested from untreated seedlings for tissue-specific expression analysis of the MsANN22-interacting proteins. Samples were immediately flash-frozen in liquid nitrogen and stored at −80□ until RNA extraction.

Total RNA was extracted from the collected samples using the HiPure HP Plant RNA Kit (Magen, Guangzhou, China) and reverse-transcribed into cDNA using the HiScript III RT SuperMix kit (Vazyme, Nanjing, China). qRT-PCR was performed using gene-specific primers listed in Supplementary Table S2. *MsACTIN* was used as the reference gene (Dong *et al*., 2021), and its expression stability under PEG, ABA, and MeJA treatments was confirmed in preliminary experiments. Each biological sample was analyzed using three technical replicates. Relative expression levels were calculated using the 2^−ΔΔCt method (Livak and Schmittgen, 2001). Significant differences were assessed by one-way ANOVA followed by Duncan’s multiple range test (*P* < 0.05).

### 2.7 Subcellular Localization Analysis

The *MsANN22* coding sequence (CDS), excluding the stop codon, was cloned into the pCAMBIA1300 vector via the XbaI and KpnI restriction sites to generate pCAMBIA1300-35S-MsANN22-eGFP. The primers are listed in Supplementary Table S2. The *MsANN51* and *MsANN72* CDSs were cloned using the same strategy to generate pCAMBIA1300-35S-MsANN51-eGFP and pCAMBIA1300-35S-MsANN72-eGFP. Positive clones were verified by Sanger sequencing and transformed into *Agrobacterium tumefaciens* strain GV3101 (Weidi, Shanghai, China). Positive *Agrobacterium* cells were cultured with shaking in LB medium containing the appropriate antibiotics until the OD□□□ reached 0.8. Cells were collected by centrifugation and resuspended to an OD□□□ of 0.6–0.7 in infiltration buffer containing 10 mM MES, 10 mM MgCl□, and 200 μM acetosyringone. The suspension carrying MsANN22-eGFP was mixed at a 1:1 ratio with a suspension carrying the cytoplasmic marker pCAMBIA1300-35S-mCherry-NOS (Point, Xi’an, China) and incubated at room temperature for 2 h. The mixture was infiltrated into the abaxial side of *Nicotiana benthamiana* leaves. Plants were maintained in the dark for 24 h and then transferred to normal light conditions for 36 h to allow accumulation of the fusion protein. Fluorescence signals were observed using a laser scanning confocal microscope (Nikon, Tokyo, Japan).

### 2.8 Cloning of *MsANN*22 and Generation of Overexpression Plants

Total RNA was extracted from alfalfa leaves and reverse-transcribed into cDNA, which was used as the template for PCR amplification of the target fragment. Amplification products were examined by agarose gel electrophoresis, and the target fragment was recovered and cloned. The resulting construct was propagated in *Escherichia coli* DH5α competent cells (Weidi, Shanghai, China), and positive single colonies were verified by sequencing. The sequence-verified overexpression vector pCAMBIA1300-35S-MsANN22-eGFP was transformed into *A. tumefaciens* GV3101 competent cells to generate the engineered strain. Alfalfa hypocotyls were used as explants. Positive *Agrobacterium* cells were resuspended in MS liquid medium to an OD□□□ of 0.4–0.6. After 10 min of inoculation, explants were co-cultivated for 2 days and transferred to callus induction medium containing 30 mg/L hygromycin for selection. Differentiated adventitious shoots were transferred to rooting medium, and positive transgenic plants were identified by genomic PCR.

*Arabidopsis* plants were transformed using the floral dip method (Ali *et al*., 2022). Harvested seeds were sown on MS selection medium containing 30 mg/L hygromycin to obtain resistant T□ transgenic lines. Lines with high target-gene expression were identified by qRT-PCR and propagated. Successive generations were screened until homozygous T□ lines were obtained for subsequent experiments.

### 2.9 Drought Treatment of Transgenic Plants and Measurement of Physiological Indicators

To investigate the role of *MsANN22* in stress tolerance, three alfalfa overexpression lines (OE-1, OE-17, and OE-22) and two *Arabidopsis* overexpression lines (OE-3 and OE-6) were subjected to stress assays at the mature-plant and germination stages.

For drought treatment, transgenic and wild-type plants with uniform growth were cultivated under normal conditions for 30 days. Watering was then withheld for 16 days to impose drought stress, followed by rewatering. Plants were photographed before treatment (day 0) and at 8 and 16 days of water withdrawal, as well as after rewatering, for phenotypic assessment. For the *Arabidopsis* seed germination assay, wild-type and overexpression seeds were surface-sterilized and placed on half-strength MS medium supplemented with 0, 150, or 200 mM mannitol. Germination rates were recorded daily. For physiological and biochemical measurements, leaf tissue (0.1 g) was homogenized in 1 mL of extraction solution on ice. The homogenate was centrifuged at 12,000 rpm for 10 min at 4□, and the supernatant was retained for analysis. Superoxide dismutase (SOD), peroxidase (POD), and catalase (CAT) activities, as well as malondialdehyde (MDA) content, were measured using assay kits (Grace Biotechnology, Jiangsu, China). Chlorophyll content was determined spectrophotometrically. Fresh leaves (2 g) were ground with quartz sand, calcium carbonate, and 95% ethanol. An additional 10 mL of 95% ethanol was added, and the extract was incubated overnight until the tissue was bleached. Absorbance of the extract was measured at 663 and 645 nm, and the total chlorophyll content was calculated according to Arnon (Arnon, 1949). Histochemical staining with nitroblue tetrazolium (NBT) and 3,3′-diaminobenzidine (DAB) was performed to visualize O□•□ and H□O□ accumulation in leaves, respectively (Zhao *et al*., 2026).

### 2.10 Construction of the Bait Vector and Autoactivation Assay

The *MsANN22* CDS was amplified by PCR and cloned into the pGBKT7 vector. The resulting plasmid was propagated in *E. coli* to obtain the bait plasmid BD-MsANN22. pGBKT7-53+pGADT7-T and pGBKT7-Lam+pGADT7-T were used as the positive and negative controls, respectively. For the autoactivation assay, the recombinant bait vector and control plasmid combinations were transformed into Y2HGold competent yeast cells and plated on quadruple-dropout medium (QDO; SD/−Trp/−Leu/−His/−Ade). Plates were incubated inverted at 30□ for 3–4 days. Colony growth was examined to determine whether the bait protein exhibited autoactivation activity. The primers are listed in Supplementary Table S2.

### 2.11 Screening of an Alfalfa cDNA Library for MsANN22-Interacting Proteins

The bait plasmid BD-MsANN22 was transformed into yeast strain Y2HGold. A positive single colony was selected and expanded in liquid medium to prepare competent cells carrying the bait plasmid. An alfalfa Y2H cDNA library was obtained from Wuhan Tianwen Biotechnology Co., Ltd. (Wuhan, China). A 30-μL aliquot of the Y2H cDNA library plasmid and 60 μL of pre-denatured salmon sperm DNA (carrier DNA) were co-transformed into these competent cells using the PEG/LiAc method. PEG and LiAc solutions were added sequentially, and the mixture was incubated in a 30 □ water bath for 45 min. After adding 20 μL of DMSO, the mixture was heat-shocked at 42 □ for 20 min. YPDA liquid medium was then added, and the cells were recovered by shaking at 30 □. A 100-μL aliquot of the recovered suspension was plated on double-dropout medium (DDO; SD/−Trp/−Leu) and incubated inverted at 30 □. Colony formation on DDO medium indicated recovery of cotransformants; interaction-dependent reporter activation was assessed in the subsequent QDO/X-α-Gal screen.

Positive single colonies on the DDO plates were picked with sterile pipette tips and spot-inoculated onto QDO/X-α-Gal selective medium (SD/−Trp/−Leu/−His/−Ade/X-α-Gal). The plates were incubated under the same conditions, and blue colonies were retained as candidate positive clones. Blue colonies were propagated in liquid medium. Plasmids were extracted, amplified by PCR using universal primers (Supplementary Table S2), and sequenced. The resulting sequences were analyzed using NCBI BLAST to identify candidate MsANN22-interacting proteins.

### 2.12 Luciferase Complementation Assay (LCA)

The CDSs of *MsANN22*, *MsIAA*, and *MsPR-10* were cloned into pCAMBIA1300-nLUC and pCAMBIA1300-cLUC by homologous recombination to generate fusion expression vectors. Recombinant and empty vectors were separately transformed into *A. tumefaciens* GV3101 competent cells. Bacterial suspensions were adjusted to an OD□□□ of 0.6 and infiltrated into the abaxial side of leaves from 6–8-week-old *N. benthamiana* plants for transient expression. The experimental combinations were MsANN22-nLUC+MsIAA-cLUC and MsANN22-nLUC+MsPR-10-cLUC. The control combinations were MsANN22-nLUC+cLUC, nLUC+MsIAA-cLUC, nLUC+MsPR-10-cLUC, and nLUC+cLUC. After infiltration, plants were incubated in the dark for 48 h. Potassium D-luciferin solution was then applied to the infiltration sites, and bioluminescence signals were visualized and recorded using an in vivo imaging system.

## 3. Results

### 3.1 Identification of the *MsANN* Gene Family in Alfalfa

Based on BLASTP searches, HMM analysis, and validation of the conserved annexin domain (PF00191), 73 annexin genes were identified in the alfalfa genome. According to their chromosomal positions, these genes were systematically designated *MsANN01* to *MsANN73*.

Chromosomal distribution analysis showed that 67 of the 73 *MsANN* genes were unevenly distributed across 28 chromosomes (Supplementary Fig. S1), with none assigned to the chr7 homeologous group. Five chromosomes (chr6.4, chr8.1, chr8.2, chr8.3, and chr8.4) harbored the largest number of *MsANN* genes (five each), followed by chr3.3 (four genes); chr3.2, chr6.1, chr6.2, and chr6.3 (three genes each); eight chromosomes (chr1.1, chr1.2, chr3.1, chr3.4, and chr4.1–chr4.4) (two genes each); and ten chromosomes (chr1.3, chr1.4, chr2.1–chr2.4, and chr5.1–chr5.4) (one gene each). The remaining six genes (*MsANN68*–*MsANN73*) were located on unanchored sequences in the genome assembly.

The physicochemical properties of the 73 MsANN proteins were analyzed (Supplementary Table S1). Protein lengths ranged from 72 (MsANN35 and MsANN41) to 327 amino acids (MsANN36, MsANN37, and MsANN42), with molecular weights of 8.25–37.43 kDa and isoelectric points of 5.39–9.37. The instability index ranged from 23.96 to 51.45 (mean 38.18), and the mean aliphatic index was 92.37, suggesting that most MsANN proteins are relatively stable and thermostable. All MsANN proteins were hydrophilic according to GRAVY analysis. Subcellular localization predictions indicated that MsANN proteins were primarily distributed in the cytoplasm, chloroplasts, nucleus, and cytoskeleton, reflecting diverse subcellular localization patterns within the family.

### 3.2 Phylogenetic Analysis of *MsANN* Genes in Alfalfa

To investigate the evolutionary relationships among plant annexins, a total of 137 ANN protein sequences, comprising the 73 MsANNs from alfalfa and the ANN members identified in *Arabidopsis*, soybean, rice, and *M. truncatula* as described in Section 2.2, were used for phylogenetic analysis. A phylogenetic tree was constructed using the ML method with 1,000 bootstrap replicates (Fig. 1). Based on their phylogenetic relationships, the ANN proteins were classified into four major groups (Groups 1–4).

**Fig. 1.**
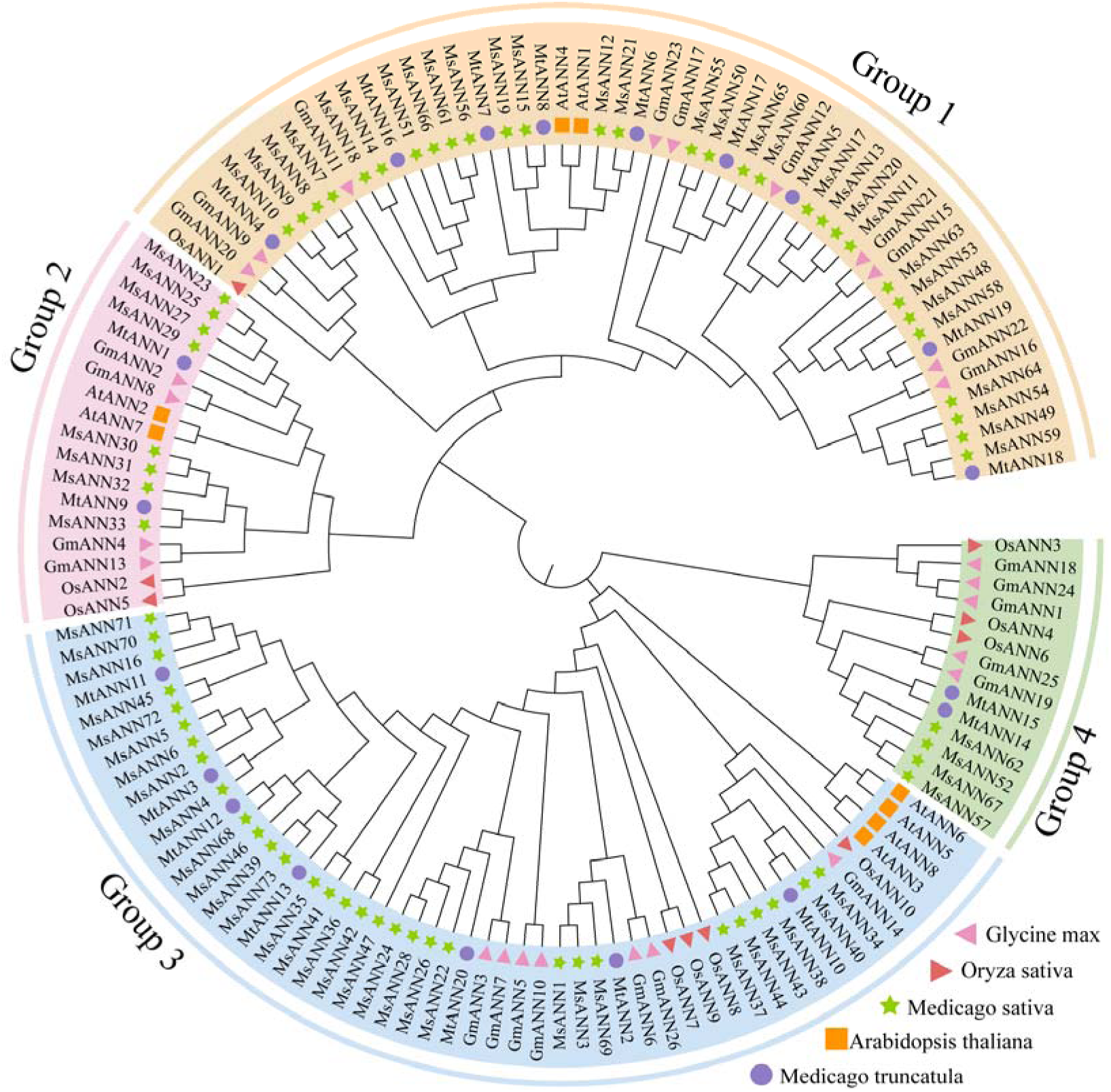
Phylogenetic analysis of annexin (ANN) proteins from *M. sativa*, *M. truncatula*, *O. sativa*, *G. max*, and *A. thaliana*. The tree was constructed using the maximum likelihood method with 1,000 bootstrap replicates. Symbols indicate species: MsANN (alfalfa, green stars), AtANN (*A. thaliana*, yellow squares), MtANN (*M. truncatula*, purple circles), GmANN (soybean, pink triangles), and OsANN (rice, red triangles).

Groups 1 and 2 contained ANN members from both dicotyledonous plants (*Arabidopsis*) and monocotyledonous plants (rice), and these members formed mixed clades with MtANNs and GmANNs. This distribution suggests that the two groups originated before the divergence of monocots and dicots, although this topology alone does not establish functional conservation. Group 3 contained the largest number of *MsANN* members (31). Within this group, MsANNs formed multiple mixed subclades with MtANNs, GmANNs, and selected AtANNs, indicating a high degree of evolutionary conservation. Group 4 comprised MsANN52, MsANN57, MsANN62, and MsANN67 but contained no AtANN members, a pattern consistent with lineage-specific diversification in legumes. Overall, MsANN members were distributed across all four phylogenetic groups and showed close evolutionary relationships with ANNs from legumes, *Arabidopsis*, and rice. Differences in clade composition suggest potential functional divergence.

### 3.3 Synteny and Gene Duplication Analysis of *MsANN* Genes

To identify duplication events within the *MsANN* family, intraspecific synteny analysis was performed for all 73 genes. Sixty-seven *MsANN* genes were unevenly distributed across all chromosomes except chr7. A total of 75 intragenomic collinear gene pairs were identified, with the greatest number involving chromosomes in the chr8 group (19 pairs), followed by the chr6 group (14 pairs) (Fig. 2). All collinear pairs were classified as segmental or whole-genome-duplication-associated pairs, whereas no tandem duplications were detected. These results suggest that large-scale duplication contributed substantially to *MsANN* family expansion. Ka/Ks ratios of all duplicated pairs were below 1 (Supplementary Table S3), indicating purifying selection on the *MsANN* family. Interspecific synteny analysis between alfalfa and the four other species (Supplementary Fig. S2) identified a total of 43, 31, 11, and 5 syntenic gene pairs between alfalfa and *M. truncatula*, soybean, rice, and *Arabidopsis*, respectively—more with dicots than with rice.

**Fig. 2.**
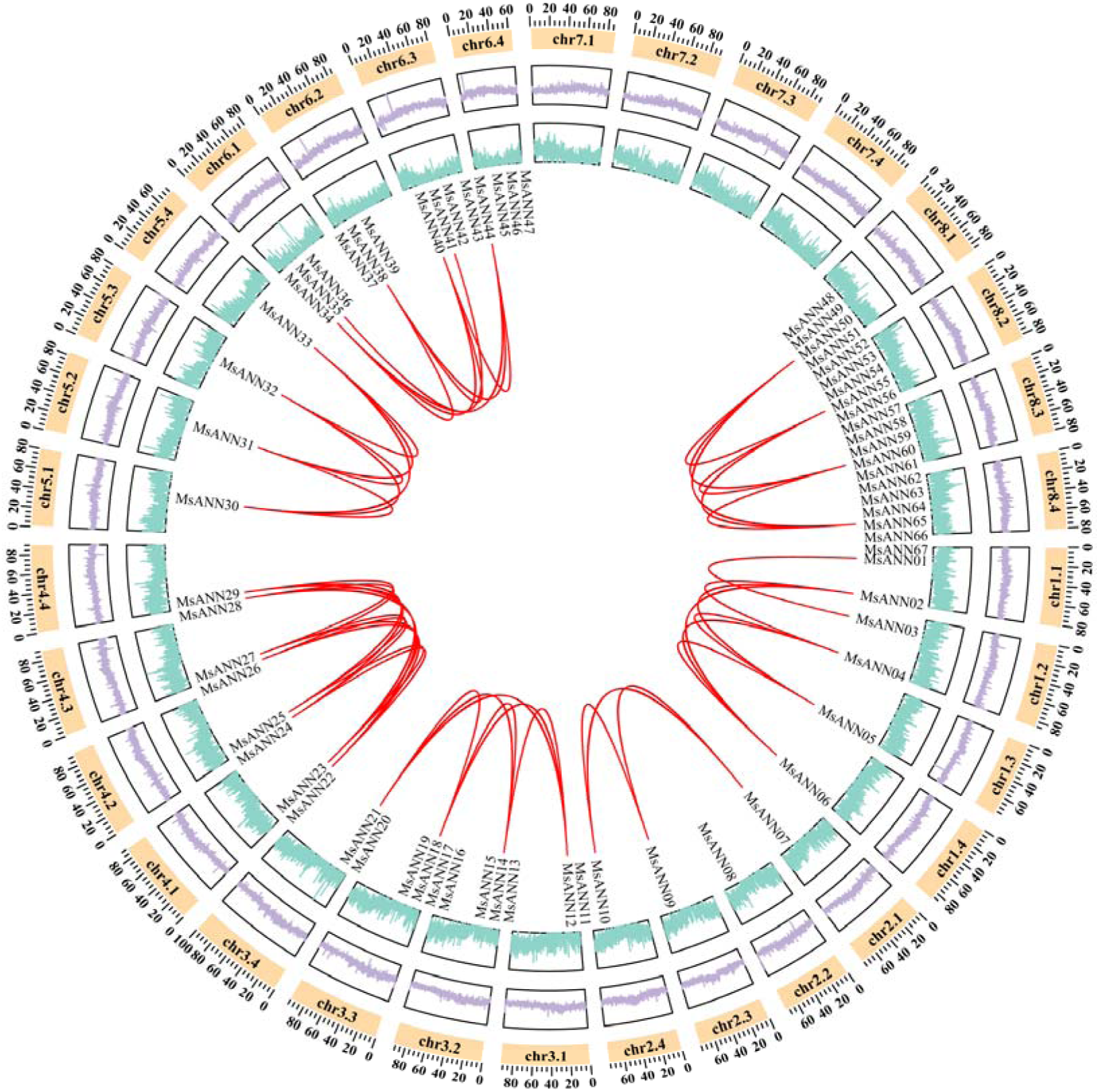
Collinearity analysis of *MsANN* genes in *Medicago sativa.* Red lines indicate intraspecific collinear gene pairs derived from segmental duplication in the *MsANN* family. Chromosome names are labeled in the inner circle, and lengths are labeled in the outer circle. Green represents gene density, and purple represents GC content.

### 3.4 Conserved Motifs, Domains, and Gene Structures of the *MsANN* Gene Family

An intraspecific ML tree of the 73 MsANN proteins resolved five Subgroups (Subgroups I–V) (Supplementary Fig. S3), which were broadly concordant with the cross-species classification: Subgroup I spanned Groups 1 and 4, Subgroup II nested within Group 1, and Subgroups III–V corresponded largely to Group 3. Conserved motif analysis of the 73 MsANN proteins using MEME identified 10 motifs (Supplementary Fig. S3). Members within the same Subgroup exhibited highly similar motif compositions and arrangements, whereas substantial differences were observed among Subgroups. Most Subgroup I members contained Motifs 1–5 and 7–9, and Motifs 1 and 3 occurred in nearly all members, suggesting that they are core elements important for MsANN function. In contrast, Subgroup V showed the greatest motif diversity, with pronounced variation among its members. Several members (*MsANN02*, *MsANN04*, *MsANN05*, *MsANN06*, *MsANN45*, *MsANN71*, and *MsANN72*) contained the complete set of Motifs 1–8, whereas a small number, including *MsANN35* and *MsANN41*, retained only some core motifs.

Conserved domain analysis detected the annexin domain (PF00191) in all MsANN proteins, with the highest conservation in Subgroups I–III. Notably, several members in Subgroups I and V (*MsANN16*, *MsANN35*, *MsANN36*, *MsANN41*, and *MsANN42*) encoded markedly shorter proteins containing only a single annexin domain, in contrast to the multiple tandem repeats (typically four) characteristic of canonical annexins, indicating domain divergence within these Subgroups. A similar pattern was observed for the ANX domain, which was evenly distributed in Subgroups I–III but varied considerably in Subgroups IV and V.

*MsANN* genes contained one to six exons, with approximately 84.9% containing six. Gene structure was most variable in Subgroup V, which included both the shortest genes (*MsANN35* and *MsANN41*, one exon each) and six-exon members such as *MsANN02* and *MsANN06*. Gene length varied most widely in Subgroup I, ranging from approximately 2,000 bp (*MsANN15*) to over 20,000 bp (*MsANN19*).

### 3.5 Analysis of Cis-Acting Elements in *MsANN* Promoters

To characterize the transcriptional regulation of *MsANN* genes, cis-acting elements in the 2,000-bp promoter regions upstream of the start codons of all 73 genes were analyzed. Thirty-nine types of regulatory elements were identified and classified into five categories: phytohormone-responsive, abiotic stress-responsive, light-responsive, growth and development-related, and transcription factor-binding elements (Fig. 3).

**Fig. 3.**
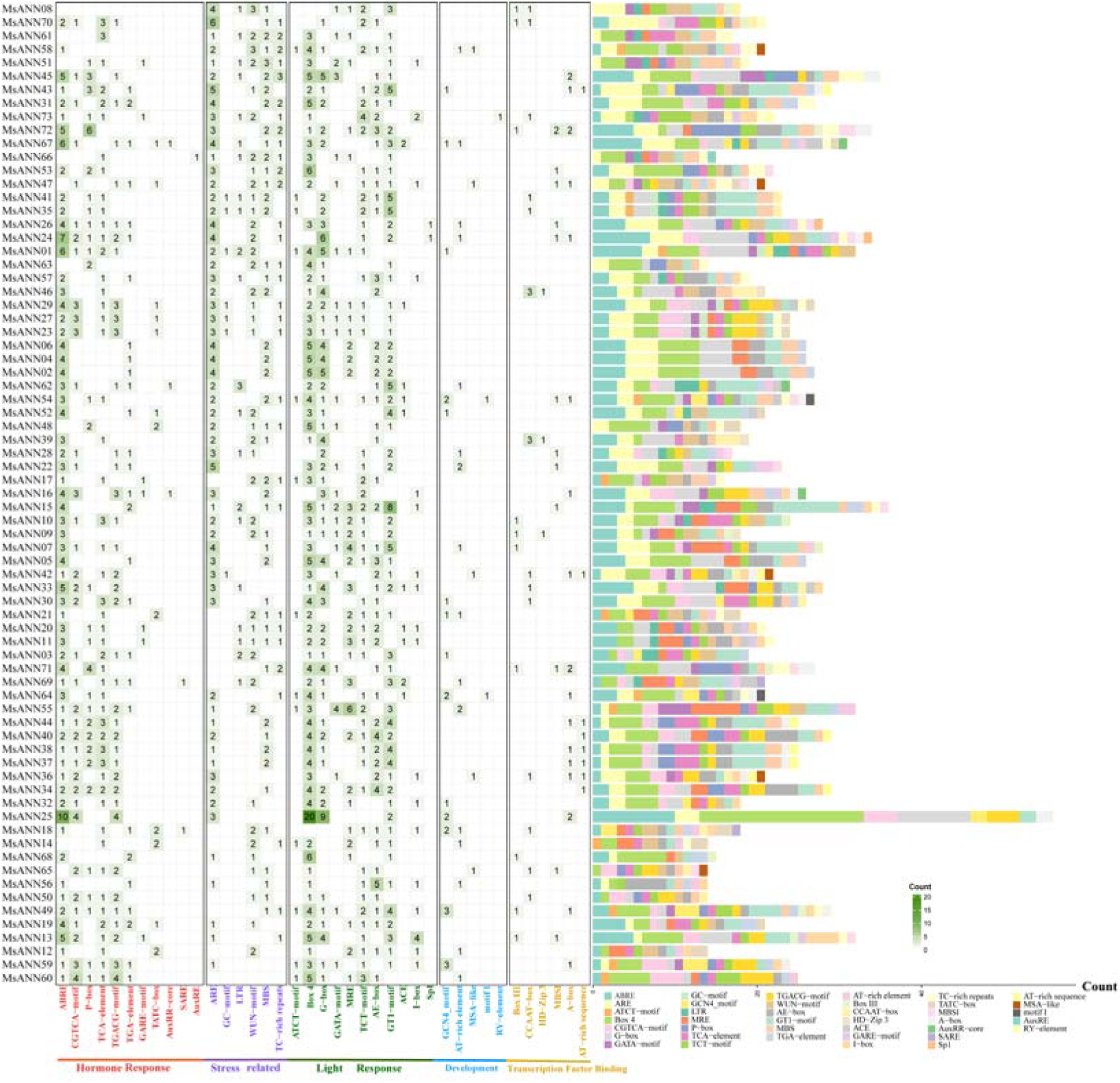
Classification and statistics of cis-acting elements in the promoter regions of *MsANN* family genes. The cis-acting element heatmap is divided into five categories. The numbers in the heatmap represent the number of cis-acting elements, and darker-colored blocks indicate a greater number of cis-acting elements. The right panel summarizes the number of cis-acting elements in the promoters. Different colors represent different categories.

Across the promoter set, *MsANN* promoters contained diverse hormone-responsive elements. The promoter of *MsANN08* was the only one lacking an ABRE; detected categories included ABA-, salicylic acid-, MeJA-, gibberellin-, and auxin-responsive elements. ABRE, CGTCA-motif, and TGACG-motif were the most widely distributed elements and were present in most *MsANN* promoters. These findings suggest that these elements may mediate hormone-dependent stress responses and coordinate plant adaptation to adverse environments.

Among the abiotic stress-responsive elements, most *MsANN* promoters contained anaerobic induction elements (ARE), low-temperature-responsive elements (LTR), drought-inducible MYB-binding sites (MBS), and wound-responsive elements (WUN-motif). In addition, TC-rich repeats, which are representative defense- and stress-responsive elements, were detected in more than 30 *MsANN* promoters. These results suggest that *MsANN* genes may contribute to transcriptional responses to oxygen deficiency, low temperature, drought, and mechanical injury.

Light-responsive elements constituted the most abundant category. The *MsANN25* and *MsANN54* promoters contained 13 and 17 light-responsive elements, respectively, indicating substantial potential for light-mediated regulation. The widespread distribution of these elements suggests that light signals may regulate *MsANN* gene expression. Overall, the composition and distribution of cis-acting elements in *MsANN* promoters were highly diverse, providing insights into the transcriptional regulation of these genes and their potential roles in plant growth, development, and stress responses.

### 3.6 Expression Patterns of *MsANN* Genes under Osmotic and Hormone Treatments

Based on the transcriptome data (Supplementary Fig. S4), six representative *MsANN* genes that responded significantly to drought stress or hormone treatments were selected for qRT-PCR analysis (Fig. 4). Alfalfa seedlings were treated with 15% PEG 6000, 50 μM MeJA, or 100 μM ABA, and the relative expression levels of *MsANN22*, *MsANN32*, *MsANN51*, *MsANN56*, *MsANN61*, and *MsANN72* were quantified in leaves and roots at multiple time points.

**Fig. 4.**
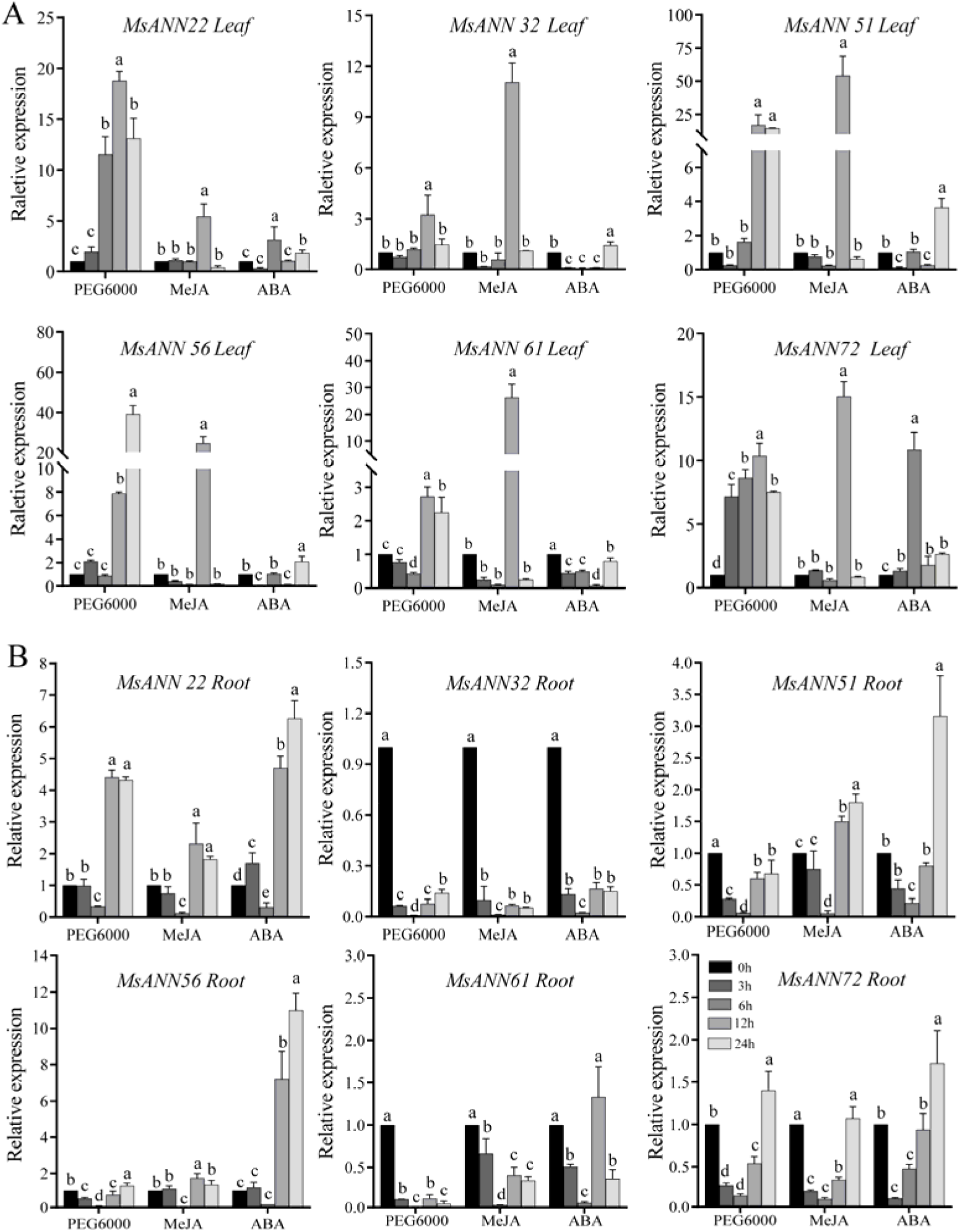
Expression profiles of selected *MsANN* genes in leaves and roots of alfalfa under ABA, MeJA and PEG 6000 treatments. (A) Leaves. (B) Roots. Relative transcript abundance was normalized to [reference gene] and is expressed relative to [0 h or untreated control], set to 1. Data are means ± SD of three independent biological replicates (n = 3); technical replicates were averaged within each biological replicate and were not treated as independent n. Statistical significance was assessed separately for each gene, tissue and treatment by one-way ANOVA followed by Duncan’s multiple range test. Different letters indicate significant differences among time points within the same tissue and treatment (*P* < 0.05). ABA, abscisic acid; MeJA, methyl jasmonate; PEG, polyethylene glycol.

Under PEG 6000-induced osmotic stress, *MsANN22* expression in leaves peaked at 12 h, whereas its expression in roots was continuously upregulated. The contrasting profiles indicate that *MsANN22* has tissue-dependent transcriptional responses. *MsANN51* and *MsANN61* were strongly induced by MeJA in leaves, with peak expression levels reaching 30-to 50-fold, but showed only weak responses in roots. Thus, these genes were preferentially responsive to MeJA in leaves under the conditions tested.

Overall, *MsANN* genes in leaves responded more strongly to MeJA than to ABA, with *MsANN51*, *MsANN61*, and *MsANN56* showing particularly high expression peaks under MeJA treatment. In roots, ABA induction was more pronounced at 24 h; *MsANN22*, *MsANN51*, *MsANN56*, *MsANN61*, and *MsANN72* all reached their highest expression levels at this time point. In these data, MeJA elicited stronger induction of several *MsANN* genes in leaves, while roots showed their strongest induction at the late ABA time point.

### 3.7 Subcellular Localization Analysis

To determine the subcellular locations of key ANN proteins, *MsANN22*, *MsANN51*, and *MsANN72* were selected based on their expression profiles. These genes showed strong transcriptional responses to PEG 6000 or hormone treatments (Fig. 4). Recombinant vectors expressing MsANN22-eGFP, MsANN51-eGFP, and MsANN72-eGFP under the control of the 35S promoter were constructed, with the 35S::eGFP vector used as the control (Fig. 5). Confocal laser scanning microscopy showed that the green fluorescence signals of all three fusion proteins strongly overlapped with the red fluorescence signal of the cytoplasmic marker mCherry, indicating that they are predominantly cytoplasmic under the normal growth conditions used in this study, although Ca²□-dependent membrane translocation cannot be excluded.

**Fig. 5.**
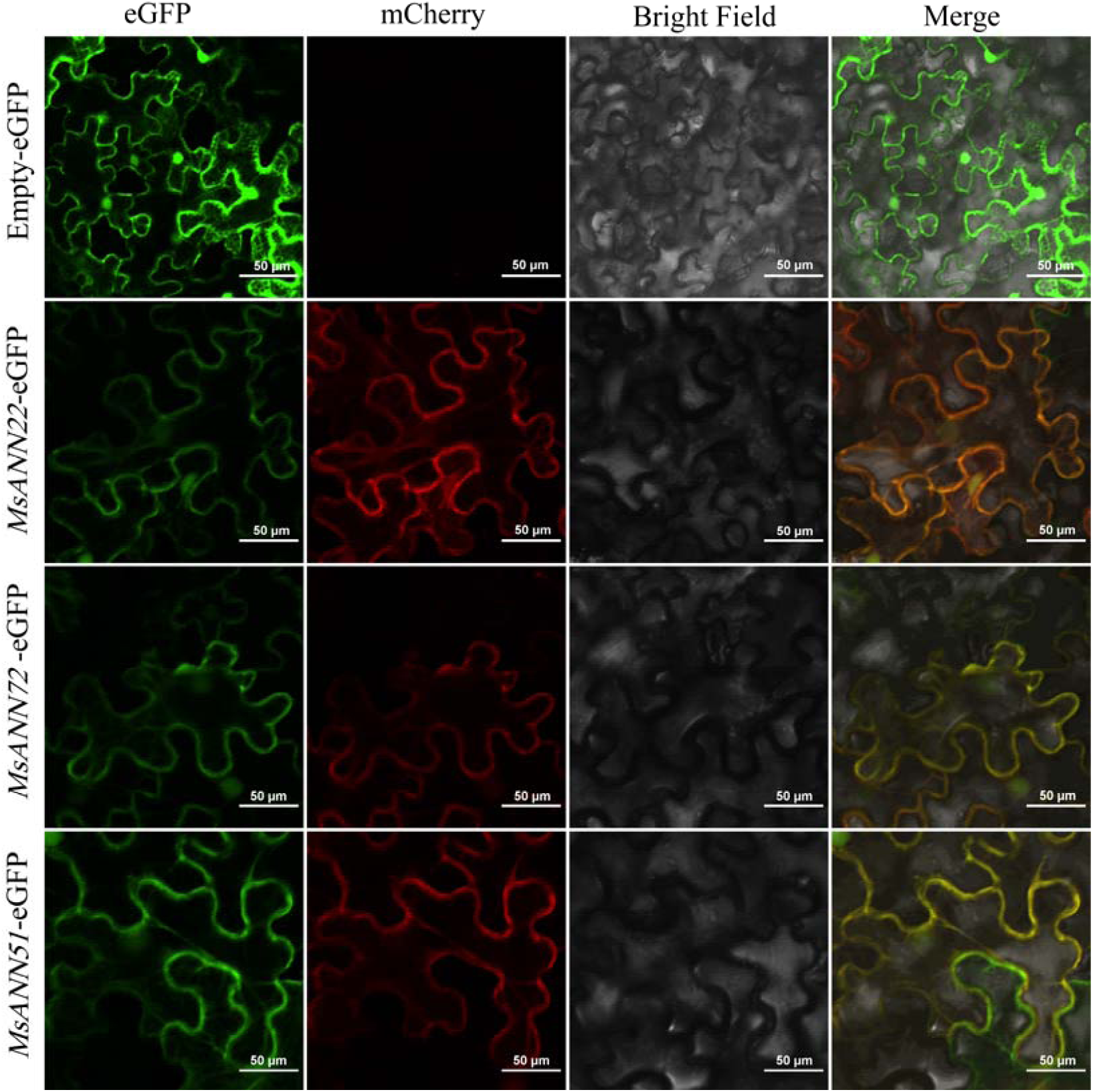
Subcellular localization of MsANN proteins in tobacco epidermal cells. eGFP: green fluorescent protein; mCherry: cytoplasmic marker; Bright: bright field; Merged: overlay of eGFP, mCherry, and bright field. Scale bar = 50 μm.

### 3.8 Overexpression of *MsANN22* Enhances Drought Tolerance in Alfalfa

To investigate the function of *MsANN22*under drought stress, pCAMBIA1300-35S-MsANN22-eGFP was introduced into alfalfa via Agrobacterium-mediated transformation, yielding three high-expression lines (OE-1, OE-17, and OE-22) for functional validation. After 16 days of water withdrawal, WT plants exhibited severe wilting and dehydration, whereas the overexpression lines maintained better growth status and leaf color (Fig. 6A); transgene expression was confirmed by PCR and qRT-PCR analyses (Fig. 6B). Following drought stress, the overexpression lines showed significantly higher leaf chlorophyll content (Fig. 6C) and only 0.8-fold the MDA content of WT plants (Fig. 6D), together with 1.38- and 1.70-fold SOD and POD activities and significantly higher CAT activity (*P* < 0.05) (Fig. 6E–G). Consistently, DAB and NBT staining revealed visibly lower H□O□ and O□•□ accumulation in the overexpression lines than in WT plants (Fig. 6H).

**Fig. 6.**
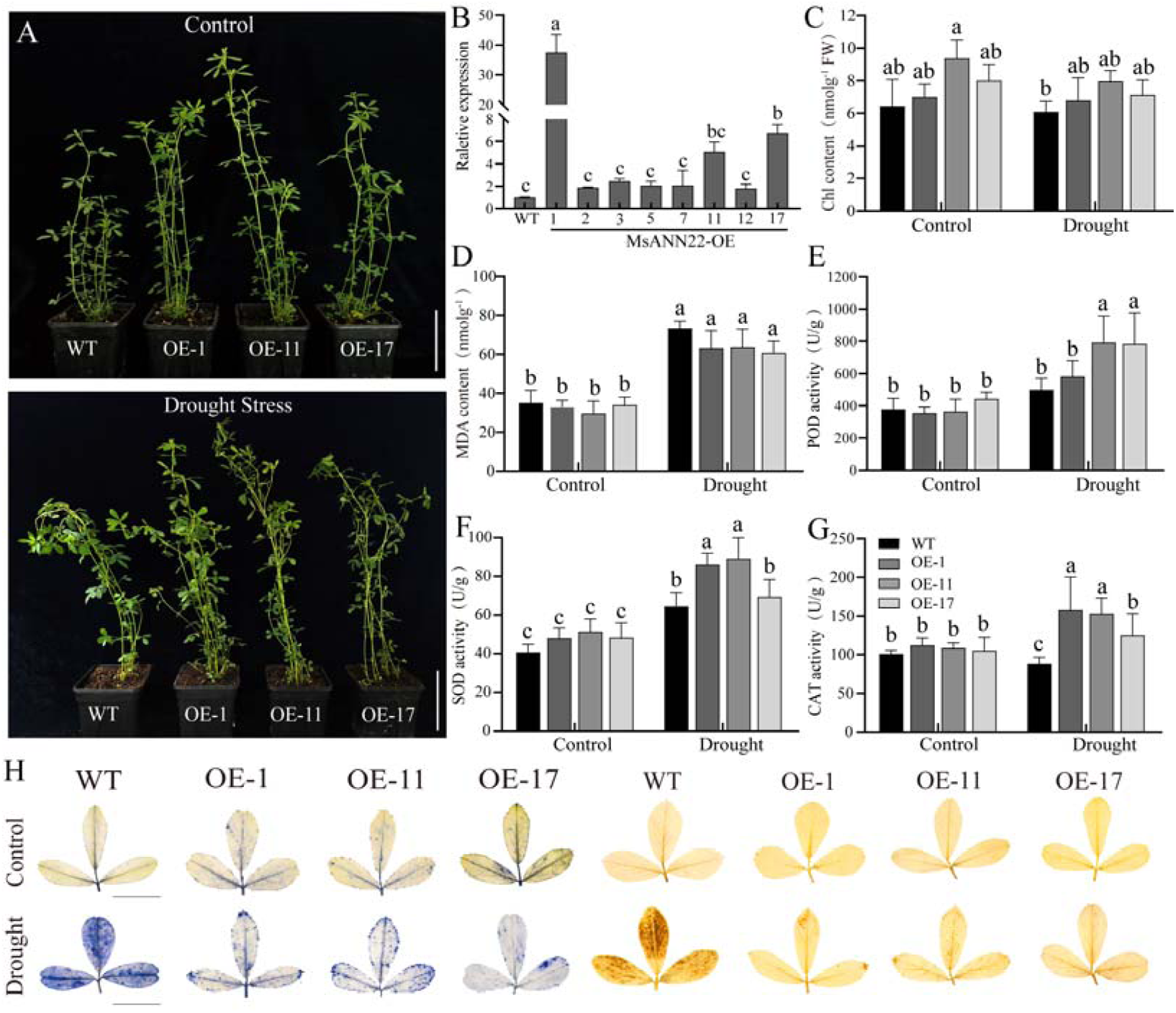
Phenotypic analysis and physiological parameter measurement of WT and *MsANN22*-overexpressing lines under drought stress. (A) Plant phenotype of wild-type (WT) and MsANN22-overexpressing (OE) alfalfa before and after drought stress. Scale bar = 10 cm. (B) Expression levels of *MsANN22* in transgenic lines. (C–G) Chlorophyll content (C), malondialdehyde (MDA) content (D), and activities of peroxidase (POD, E), superoxide dismutase (SOD, F) and catalase (CAT, G) in WT and OE lines before and after drought stress. (H) NBT and DAB staining of leaves from WT and OE plants before and after drought stress to visualize superoxide and hydrogen peroxide accumulation, respectively. Scale bar = 2 cm. Data are shown as means ± SD of three independent biological replicates (n = 3). Statistical analysis was performed by one-way ANOVA followed by Duncan’s multiple range test. Different letters indicate significant differences (*P* < 0.05).

### 3.9 Overexpression of *MsANN22* Enhances Drought Tolerance in *Arabidopsis*

To further validate the function of *MsANN22* under drought stress, two transgenic *Arabidopsis* lines (OE-3 and OE-6) and WT plants were subjected to drought treatment. After water was withheld from four-week-old plants for 8 days, WT plants exhibited pronounced leaf yellowing, whereas the transgenic lines showed no obvious changes. After 16 days of drought stress, WT plants exhibited more severe wilting than the transgenic lines. Following rewatering, OE-6 showed greater visual recovery, whereas WT plants showed limited visual recovery (Fig. 7A). These observations suggest that *MsANN22* overexpression improves recovery after drought treatment. Seed germination was subsequently evaluated on half-strength MS medium containing different mannitol concentrations (Fig. 7B). Germination rates did not differ significantly between WT and the overexpression lines at 0 mM mannitol. Under 150 and 200 mM mannitol, the germination rates of OE-3 and OE-6 were higher than that of WT, and the difference was statistically significant at 200 mM. These findings indicate that *MsANN22* overexpression improves seed germination under mannitol-induced osmotic stress.

**Fig. 7.**
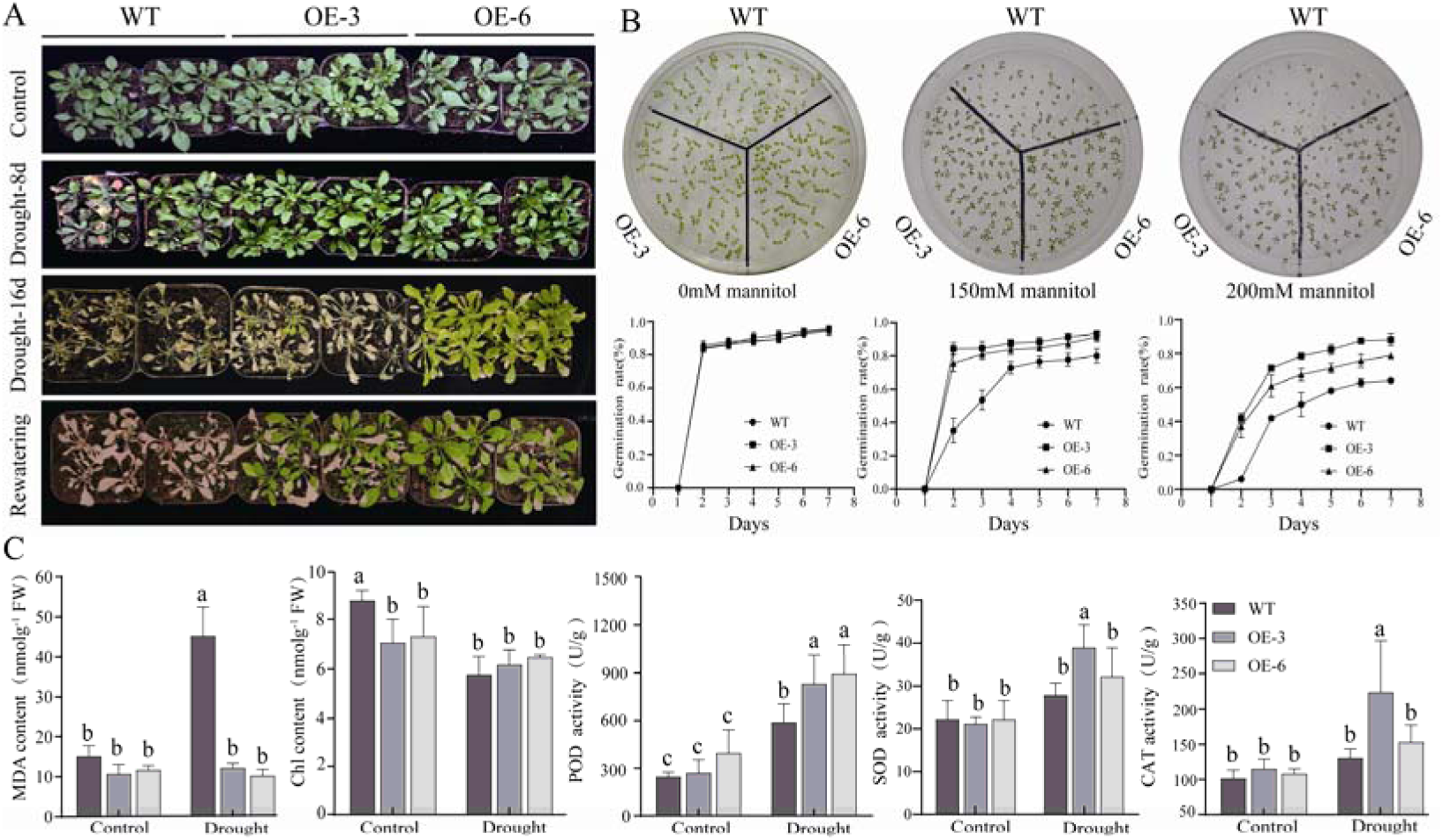
Phenotypes, seed germination rates, and physiological indices of MsANN22-overexpressing Arabidopsis under control and stress conditions. (A) Phenotypes of *MsANN22*-overexpressing (OE) Arabidopsis under normal conditions, after 8 or 16 d of drought, and after rewatering. (B) Seed germination and germination rates of wild-type (WT) and *MsANN22*-OE lines (OE-3 and OE-6) after 7 d on medium containing 0, 150, or 200 mM mannitol. (C) Malondialdehyde (MDA) content, chlorophyll content, and peroxidase (POD), superoxide dismutase (SOD), and catalase (CAT) activities in WT and *MsANN22*-OE Arabidopsis under normal conditions and drought stress. Data are shown as means ± SD of three independent biological replicates (n = 3). Statistical analysis was performed by one-way ANOVA followed by Duncan’s multiple range test. Different letters indicate significant differences (*P <* 0.05).

Under normal conditions, no significant differences in the measured physiological indicators were observed between WT and the overexpression lines. Under drought stress, however, the MDA content was significantly lower in the overexpression plants than in WT plants, whereas SOD, POD, and CAT activities and chlorophyll content were significantly higher (Fig. 7C). These results indicate that *MsANN22* overexpression is associated with improved germination under mannitol-induced osmotic stress, better chlorophyll retention, and higher antioxidant enzyme activities during soil drying.

### 3.10 Screening and Validation of MsANN22-Interacting Proteins

To identify potential molecular partners of MsANN22, a yeast two-hybrid assay was used to screen for interacting proteins. The BD-MsANN22 bait plasmid and the positive control were separately transformed into Y2HGold competent cells for autoactivation assays. The positive control (pGBKT7-53+pGADT7-T) grew normally on QDO selective medium (SD/−Trp/−Leu/−His/−Ade), whereas yeast cells carrying pGBKT7-MsANN22+pGADT7 did not grow. These results showed no detectable autoactivation by MsANN22 under the conditions tested (Supplementary Fig. S5).

Competent yeast cells carrying BD-MsANN22 were subsequently used to screen a Y2H cDNA library. Positive clones were selected on QDO/X-α-Gal medium and sequenced. BLAST analysis identified two candidate MsANN22-interacting proteins, which were designated MsIAA and MsPR-10. Bioinformatic characterization showed that MsIAA (MS.gene44001.t1), identified from the alfalfa genome in this study, is a member of the Aux/IAA transcriptional repressor family. It encodes a 253-amino-acid protein with a predicted molecular weight of 27.51 kDa and contains a typical Aux/IAA family domain. MsPR-10 (MS.gene059046.t1) encodes a 157-amino-acid protein with a predicted molecular weight of 16.64 kDa. This protein contains a typical Bet v1 domain and belongs to the pathogenesis-related protein PR-10 family. Tissue expression analysis showed that *MsIAA* was most highly expressed in roots, followed by stems, whereas *MsPR-10* was highly expressed in leaves and expressed at relatively low levels in roots and stems (Supplementary Fig. S6).

To validate the candidate interactions, BD-MsANN22 was co-transformed with AD-MsIAA or AD-MsPR-10 into yeast cells for directed pairwise Y2H assays. Both the positive control (pGBKT7-53+pGADT7-T) and the experimental combinations grew normally and activated the reporter genes, supporting the candidate interactions (Fig. 8A). These candidate interactions were further evaluated by luciferase complementation. MsANN22-nLUC was co-expressed with MsIAA-cLUC or MsPR-10-cLUC in *N. benthamiana* leaves. Strong bioluminescent signals were detected in the co-expression regions, whereas no obvious signals were observed in the empty-vector or single-construct controls (Fig. 8B). These results further supported interactions of MsANN22 with MsIAA and MsPR-10 in transiently expressed *N. benthamiana* leaves.

**Fig. 8.**
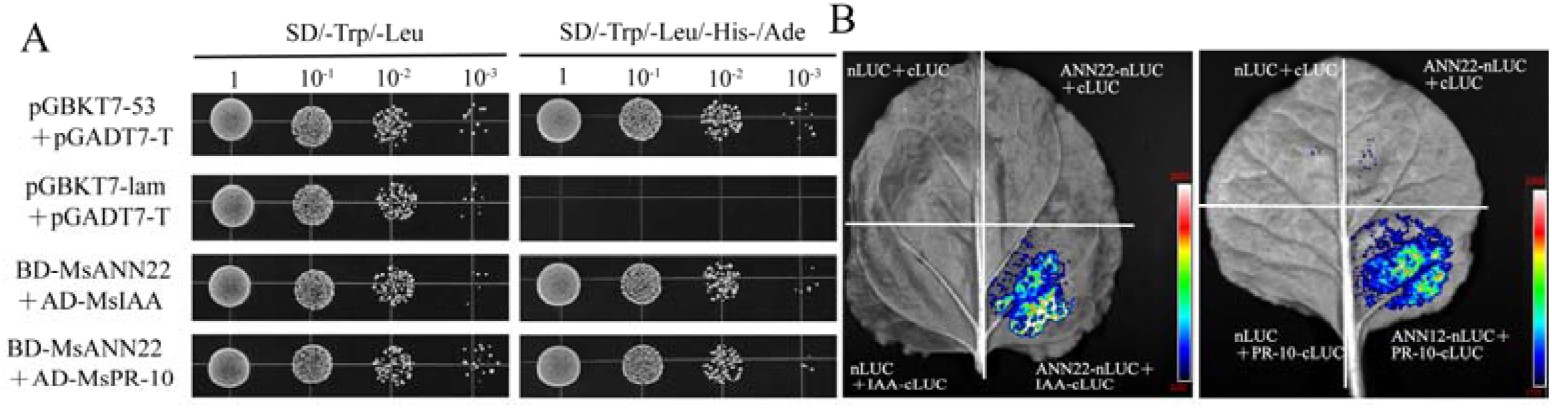
Validation of MsANN22-interacting proteins. (A) Yeast two-hybrid directed (pairwise) assay. Growth of yeast strains on SD/−Trp/−Leu/−His/−Ade medium. Positive control: pGBKT7-53+pGADT7-T; negative control: pGBKT7-MsANN22+pGADT7. Experimental groups: BD-MsANN22+AD-MsIAA; BD-MsANN22+AD-MsPR-10; (B) Luciferase complementation imaging (LCI) assay in *N. benthamiana* leaves. Bioluminescence was detected after co-expression of MsANN22-nLUC with MsIAA-cLUC or MsPR-10-cLUC. Negative controls included empty vector combinations and single-transformation combinations.

## 4. Discussion

Annexins are conserved calcium-dependent proteins that participate in plant growth and stress responses by regulating calcium signaling and maintaining membrane stability (Gerke *et al*., 2005; Laohavisit *et al*., 2010). Functional studies of annexins under abiotic stress have primarily focused on *Arabidopsis*, rice, tomato, and black cottonwood (*Populus trichocarpa*). In legumes, only a limited number of studies have been conducted in soybean and *M. truncatula* (Cantero et al., 2006; Feng *et al*., 2013; Wei et al., 2022). Molecular studies of drought tolerance in tetraploid cultivated alfalfa remain particularly scarce. Alfalfa is a perennial leguminous forage crop with high nutritional value, and drought is a major constraint on its productivity. In this study, 73 ANN family members were systematically identified in the alfalfa genome. By contrast, only 8, 26, and 20 ANN members have been reported in diploid *Arabidopsis*, soybean, and *M. truncatula*, respectively. This difference is likely associated with the autotetraploid nature of cultivated alfalfa and the whole-genome and segmental duplication events that occurred during its evolution, which collectively promoted the expansion of the ANN gene family.

Phylogenetic analysis classified 137 ANN proteins from alfalfa, *Arabidopsis*, soybean, rice, and *M. truncatula* into four groups (Fig. 1). However, when a phylogenetic tree was constructed using only alfalfa annexins and analyzed together with gene structures and conserved motifs, five Subgroups were identified. Clades III and V contained only alfalfa annexin members, although all of these genes were assigned to Group 3 in the cross-species classification. This pattern is consistent with lineage-specific diversification in the alfalfa genome. The number of ANN subfamilies varies among species. The ANN families of crape myrtle and black cottonwood contain three groups (Wei et al., 2022; Wei *et al*., 2024), whereas those of pepper, tomato, and rice contain five groups (Jami *et al*., 2012; Lu et al., 2012; Wu *et al*., 2021). Six groups have been identified in wheat (Xu *et al*., 2016). In the present phylogenetic tree, 30 MsANN proteins in Group 1 clustered with AtANN1 and AtANN4, identifying MsANN members in this Subgroup as candidates for functional comparison with AtANN1 and AtANN4.

Gene duplication is a major mechanism driving gene family expansion. In alfalfa, all 75 intragenomic collinear pairs were classified as segmental or whole-genome-duplication-associated pairs, and no tandem duplications were detected (Fig. 2). Together, these results suggest that large-scale duplication contributed substantially to *MsANN* family expansion, consistent with the expansion pattern of the annexin family in black cottonwood (Wei et al., 2022). Interspecific synteny analysis identified 43 syntenic relationships between alfalfa and *M. truncatula*, substantially more than the five syntenic relationships detected between alfalfa and *Arabidopsis* (Supplementary Fig. S2). Phylogenetic analysis further supported high sequence homology between *MsANN* and *MtANN* members, indicating a closer evolutionary relationship between alfalfa and *M. truncatula* than between alfalfa and *Arabidopsis*. Gene structure analysis can reveal important evolutionary events within gene families (Betts *et al*., 2001). Most MsANN proteins contained the conserved annexin core domain (Supplementary Fig. S3), which is associated with Ca²□-dependent phospholipid binding. In contrast, their N-terminal domains varied in sequence and length (Qiao et al., 2015). Members within the same subgroup generally shared similar gene structures and motif compositions.

The types and numbers of predicted promoter cis-elements may contribute to differential transcriptional responses to abiotic stress (Wu et al., 2021). In this study, all *MsANN* promoters except that of *MsANN08* contained ABREs (Fig. 3), suggesting potential ABA responsiveness but not, by itself, demonstrating regulation through an ABA-dependent pathway (Singh *et al*., 2015). Previous studies support this interpretation. Increased endogenous ABA levels enhance drought tolerance in rice (Li et al., 2019), whereas regulation of ABA accumulation improves salt tolerance in cotton. ABREs are highly conserved and widely distributed in ANN promoters from monocots, such as rice and wheat, and dicots, including soybean, cotton, tomato, common bean, and pepper. This conservation likely reflects broadly conserved hormone-responsive potential, although motif presence alone does not establish physiological function. In addition, *MsANN* promoters contained salicylic acid-, MeJA-, gibberellin-, and auxin-responsive elements, suggesting that these genes may integrate multiple hormone signals in response to environmental changes. The *AtANN1* promoter is enriched in the drought-responsive elements DRE and MBS, and *AtANN1* overexpression induces DREB and MYB transcription factors (Konopka-Postupolska *et al*., 2009). Consistent with this finding, MBS and ARE elements were identified in *MsANN* promoters, supporting the potential involvement of these genes in drought stress regulatory networks. Collectively, these predictions suggest that *MsANN* genes may respond to multiple hormonal and stress-related inputs, a hypothesis that requires direct regulatory assays.

To validate the transcriptome data, six representative MsANN genes responsive to drought stress or hormone treatments were selected for qRT-PCR analysis, which showed expression trends similar to those in the transcriptome data. Among the drought-responsive members, *MsANN22* was selected for detailed functional characterization because, among the six genes examined by qRT-PCR, it showed strong and sustained induction in leaves and roots under PEG 6000-induced osmotic stress, its promoter contained multiple ABA- and drought-responsive cis-elements, and it was expressed in tissues relevant to drought adaptation. Other strongly induced candidates (*MsANN51* and *MsANN61*) were predominantly induced by MeJA in a leaf-specific manner and were therefore not prioritized for transgenic functional analysis. Having confirmed the osmotic-stress-responsive expression of *MsANN22*, we next examined how its overexpression affects the physiological consequences of drought stress. Oxidative stress-induced membrane damage and loss of membrane stability are important indicators of drought-induced injury in plants. To eliminate ROS and other toxic compounds, plants increase the activities of enzymatic antioxidants, including SOD, CAT, ascorbate peroxidase (APX), POD, and glutathione reductase (GR), thereby mitigating oxidative damage (Prochazkova *et al*., 2001). Under drought stress, *MsANN22*-overexpressing *Arabidopsis* and alfalfa plants exhibited higher POD, CAT, and SOD activities and lower MDA contents than WT plants. In alfalfa, DAB staining showed lower H□O□ accumulation in the transgenic plants (Fig. 6H). These results indicate that *MsANN22* overexpression alleviates drought-induced membrane lipid peroxidation and enhances antioxidant defense capacity.

Promoter analysis showed that the *MsANN22* promoter contains predicted stress-responsive elements, including ABRE and ARE. These motifs identify ABA and redox regulation as testable hypotheses for upstream and downstream analysis of *MsANN22*. However, the hierarchical and synergistic relationships among calcium signaling, ABA signaling, and ROS scavenging remain unresolved and require direct experimental validation. In addition, although annexins are defined as Ca²□-dependent phospholipid-binding proteins and Ca²□ signaling is implicated throughout this study, the Ca²□-binding activity of *MsANN22* and its Ca²□-dependent membrane association were not directly tested; these properties remain to be demonstrated experimentally (e.g., by liposome co-sedimentation in the presence of Ca²□ or the Ca²□ chelator EGTA). It should also be noted that the present conclusion that *MsANN22* functions as an endogenous regulator of drought tolerance is based primarily on gain-of-function (overexpression) evidence; definitive loss-of-function validation—for example, via CRISPR/Cas9 knockout or RNAi/CRISPRi knockdown of *MsANN22* in alfalfa, or complementation of an *Arabidopsis* ann1/ann4 mutant—remains to be performed.

Yeast two-hybrid screening identified MsIAA and MsPR-10 as candidate MsANN22-interacting proteins, and these interactions were further supported in transiently expressed N. benthamiana leaves by the luciferase complementation assay (Fig. 8B). Aux/IAA proteins are key negative regulators of auxin signaling and typically interact with ARF transcription factors to regulate downstream gene expression (Liscum and Reed, 2002). Several Aux/IAA proteins participate directly in drought and salt stress responses (Shani *et al*., 2017), and their expression is induced by ABA and abiotic stress (Jain and Khurana, 2009). Interactions between annexins and Aux/IAA proteins have rarely been reported. The interaction between MsANN22 and MsIAA raises the hypothesis that MsANN22 could influence auxin-associated drought responses; this possibility requires functional testing. PR-10 proteins are pathogenesis-related proteins involved in plant defense against biotic and abiotic stresses, and some members possess RNase activity or antioxidant functions (Liu and Ekramoddoullah, 2006). PR-10 genes are upregulated under drought, salt, and ABA treatments in multiple species (Colditz *et al*., 2004; Hashimoto *et al*., 2004; Bahramnejad *et al*., 2010). PR-10 proteins may also enhance stress tolerance by scavenging ROS or regulating hormone signaling (Cao *et al*., 2024). Therefore, the interaction between MsANN22 and MsPR-10 raises a testable link between MsANN22 and PR-10-associated stress responses. This model remains hypothetical, and the biological functions of the MsANN22–MsIAA and MsANN22–MsPR-10 interactions, as well as their downstream signaling pathways, should be further validated using approaches such as bimolecular fluorescence complementation, co-immunoprecipitation, and genetic interaction analysis.

In summary, 73 annexin gene family members were systematically identified and characterized in alfalfa and classified into four phylogenetic groups. These genes were unevenly distributed across chromosomes and contained one to six exons. Gene structure and conserved motif analyses revealed highly conserved sequence features among members of the same Subgroup. Promoter analysis showed that *MsANN* genes were enriched in ABA-, MeJA-, and drought-responsive elements. Expression profiling showed that *MsANN22* was induced by PEG-mediated osmotic stress and ABA treatment. *MsANN22* overexpression increased antioxidant enzyme activities, reduced ROS accumulation and membrane lipid peroxidation, and was associated with improved performance in transgenic *Arabidopsis* and alfalfa under drought-related treatments. Yeast two-hybrid and luciferase complementation assays supported interactions of MsANN22 with MsIAA and MsPR-10 (Fig. 9). These findings provide a foundation for elucidating the functions of *MsANN* genes in drought stress responses and identify *MsANN22* as a candidate gene for improving drought tolerance in alfalfa. Further investigation of the downstream signaling pathways associated with these interacting proteins is required to determine the biological significance of these interactions under drought stress.

**Fig. 9.**
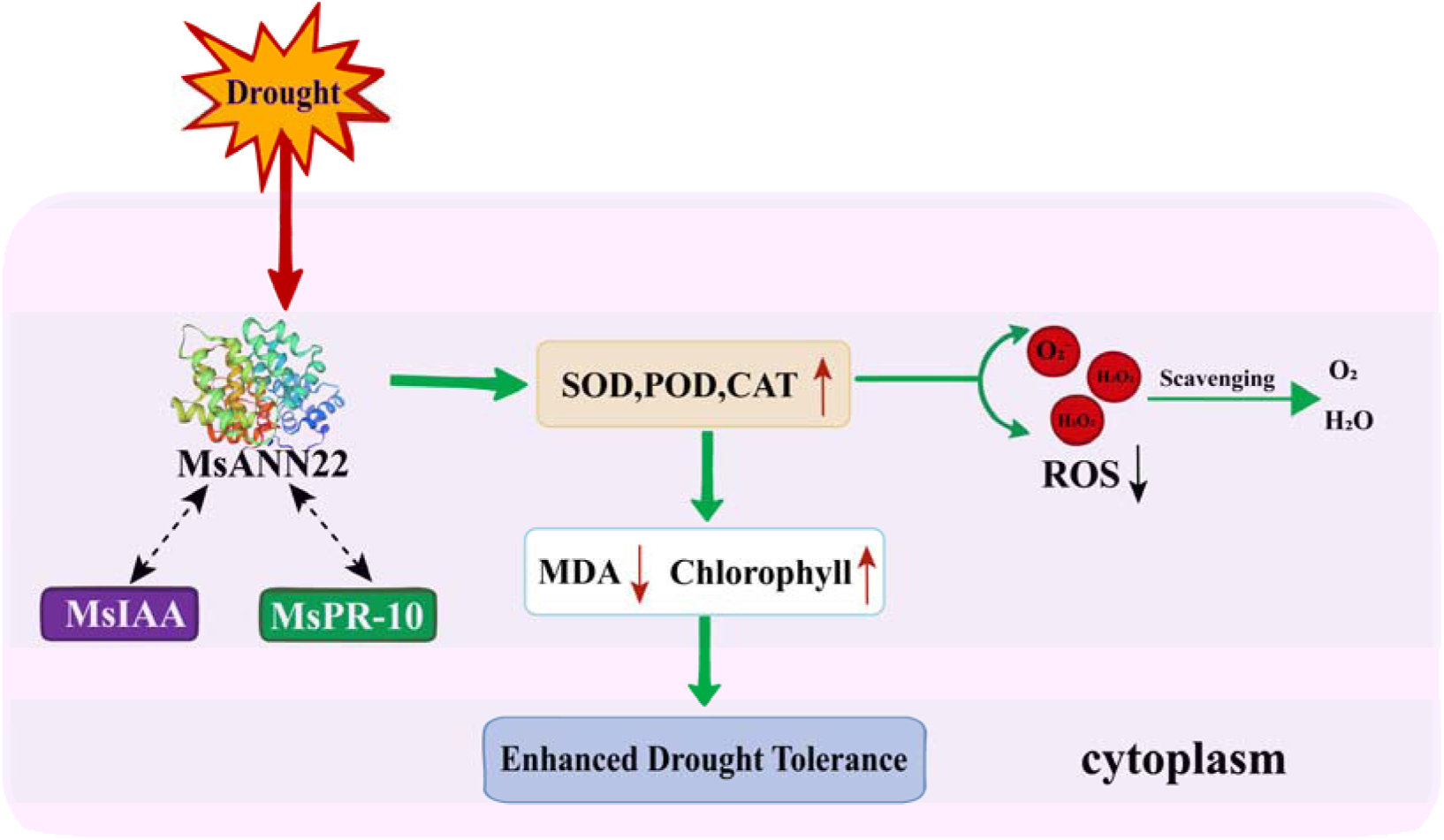
A proposed working model for MsANN22-mediated drought tolerance in alfalfa. Drought stress induces the expression of *MsANN22*, whose encoded protein is localized in the cytoplasm. *MsANN22* overexpression is associated with higher activities of antioxidant enzymes, including superoxide dismutase (SOD), peroxidase (POD), and catalase (CAT), together with lower accumulation of O • and H O under drought stress. In addition, MsANN22 shows candidate interactions with MsIAA and MsPR-10, as supported by yeast two-hybrid (Y2H) and luciferase complementation (LCA) assays. Lower ROS accumulation was accompanied by reduced malondialdehyde (MDA) content and better chlorophyll retention, consistent with reduced oxidative injury under drought stress. Solid green arrows indicate promotion, the green blunt-ended line indicates scavenging/inhibition, and dashed black arrows indicate protein–protein interactions; ↑ and ↓ indicate increased and decreased levels or activities, respectively.

## 5. Conclusion

This study performed a genome-wide identification and systematic characterization of the annexin gene family in alfalfa, identifying 73 *MsANN* members classified into four phylogenetic groups. The family has expanded mainly through whole-genome and segmental duplications, consistent with the autotetraploid origin of cultivated alfalfa. Promoter analysis showed that *MsANN* promoters are enriched in ABA- and drought-responsive cis-elements, and transcriptome profiling further identified *MsANN22* as a member strongly induced by drought and ABA. Functional assays demonstrated that *MsANN22* overexpression enhanced drought tolerance in both *Arabidopsis* and alfalfa by increasing antioxidant enzyme activities, maintaining chlorophyll content, and reducing ROS and MDA accumulation. MsANN22 also interacts with MsIAA and MsPR-10, suggesting its involvement in hormone-and defense-related pathways. These results provide a genomic resource for the alfalfa annexin family and identify *MsANN22* as a candidate gene for molecular breeding of drought-tolerant alfalfa.

## Supplementary data

The following supplementary data are available at JXB online.

Fig. S1. Chromosomal localization analysis of *MsANN* genes.

Fig. S2. Collinearity analysis of annexin genes between *Medicago sativa*, *Glycine max*, *Medicago truncatula*, *Arabidopsis thaliana* and *Oryza sativa*.

Fig. S3. Phylogenetic relationships, conserved motifs, conserved domains, and gene structures of *MsANN* geness.

Fig. S4. Expression profiling heatmap of *MsANN* genes under drought, salt, and jasmonic acid treatments.

Fig. S5. Autoactivation assay of BD-MsANN22 in the yeast two-hybrid (Y2H) system.

Fig. S6. Expression levels of *MsPR-10* and *MsIAA* in roots, stems, and leaves.

Table S1. The detail information of ANN protein in M. sativa

Table S2. Primer List

Table S3. Analysis of ANN gene pairs for KaKs in M. sativa

## Author contributions

Zhengjun Sheng: Writing – original draft, Methodology, Formal analysis, Data curation. Guangzhi Jiang: Writing – original draft, Methodology, Formal analysis. Jiayin Wang: Writing – original draft, Data curation, Validation. Huan He: Data curation, Validation. Pingping Du: Data curation, Validation. Hongfei Li: Investigation, Data curation. Zhuang Meng: Resources, Project administration, Funding acquisition. Qinglong Shao: Resources, Project administration, Funding acquisition. Hongbin Li: Writing – review & editing, Resources, Project administration, Funding acquisition. Quanliang Xie: Writing – review & editing, Conceptualization, Supervision, Project administration, Funding acquisition.

## Data availability

Data will be made available on request.

## References

Ali I, Salah K B H, Sher H, et al. 2022. Drought stress enhances the efficiency of floral dip method of Agrobacterium-mediated transformation in Arabidopsis thaliana. Brazilian Journal of Biology. 84, e259326.

Andrawis A, Solomon M. Delmer D P. 1993. Cotton fiber annexins: a potential role in the regulation of callose synthase. The Plant Journal. 3, 763–772.

Arnon D I. 1949. Copper enzymes in isolated chloroplasts. Polyphenoloxidase in Beta vulgaris. Plant Physiology. 24, 1–15.

Bahramnejad B, Goodwin P H, Zhang J, et al. 2010. A comparison of two class 10 pathogenesis-related genes from alfalfa and their activation by multiple stresses and stress-related signaling molecules. Plant Cell Reports. 29, 1235–1250.

Bailey T L, Boden M, Buske F A, et al. 2009. MEME SUITE: tools for motif discovery and searching. Nucleic Acids Research. 37, W202–208.

Baucher M, Oukouomi Lowe Y, Vandeputte O M, et al. 2011. Ntann12 annexin expression is induced by auxin in tobacco roots. Journal of Experimental Botany. 62, 4055–4065.

Betts M J, Guigó R, Agarwal P, et al. 2001. Exon structure conservation despite low sequence similarity: a relic of dramatic events in evolution? The EMBO journal. 20, 5354–5360.

Bolger A M, Lohse M. Usadel B. 2014. Trimmomatic: a flexible trimmer for Illumina sequence data. Bioinformatics. 30, 2114–2120.

Cantero A, Barthakur S, Bushart T J, et al. 2006. Expression profiling of the Arabidopsis annexin gene family during germination, de-etiolation and abiotic stress. Plant Physiology Biochemistry. 44, 13–24.

Cao J, Maitirouzi A, Feng Y, et al. 2024. Heterologous expression of Halostachys caspica pathogenesis-related protein 10 increases salt and drought resistance in transgenic Arabidopsis thaliana. Plant Molecular Biology. 115, 5.

Chen C, Chen H, Zhang Y, et al. 2020. TBtools: An Integrative Toolkit Developed for Interactive Analyses of Big Biological Data. Molecular Plant. 13, 1194–1202.

Chen K, Renaut J, Sergeant K, et al. 2013. Proteomic changes associated with freeze-thaw injury and post-thaw recovery in onion (Allium cepa L.) scales. Plant Cell Environ. 36, 892–905.

Clark G B, Morgan R O, Fernandez M P, et al. 2012. Evolutionary adaptation of plant annexins has diversified their molecular structures, interactions and functional roles. New Phytologist. 196, 695–712.

Colditz F, Nyamsuren O, Niehaus K, et al. 2004. Proteomic approach: Identification of proteins induced in roots after infection with the pathogenic oomycete. Plant Molecular Biology. 55, 109–120.

DeFalco T A, Chiasson D, Munro K, et al. 2010. Characterization of GmCaMK1, a member of a soybean calmodulin-binding receptor-like kinase family. FEBS letters. 584, 4717–4724.

Divya K, Jami S K. Kirti P B. 2010. Constitutive expression of mustard annexin, AnnBj1 enhances abiotic stress tolerance and fiber quality in cotton under stress. Plant Molecular Biology. 73, 293–308.

Dong X M, Deng H, Ma W X, et al. 2021. Genome-wide identification of the MADS-box transcription factor family in autotetraploid cultivated alfalfa and expression analysis under abiotic stress. BMC Genomics. 22, 603.

Faizan M, Ashoka H, Karabulut F, et al. 2024. Annexins: A family of calcium binding proteins with variety of roles in plant development and abiotic stress tolerance. Plant Stress. 14, 100573.

Feng Y M, Wei X K, Liao W X, et al. 2013. Molecular analysis of the annexin gene family in soybean. Biologia Plantarum. 57, 655–662.

Gao Y L, Cui Y J, Li M N, et al. 2024. Comparative proteomic discovery of salt stress response in alfalfa roots and overexpression of MsANN2 confers salt tolerance. Plant Physiology Biochemistry. 215, 109033.

Gerke V, Creutz C E. Moss S E. 2005. Annexins: linking Ca²□ signalling to membrane dynamics. Nature Reviews Molecular Cell Biology. 6, 449–461.

Gerke V. Moss S E. 2002. Annexins: From structure to function. Physiological Reviews. 82, 331–371.

Hashimoto M, Kisseleva L, Sawa S, et al. 2004. A novel rice PR10 protein, RSOsPR10, specifically induced in roots by biotic and abiotic stresses, possibly via the jasmonic acid signaling pathway. Plant and Cell Physiology. 45, 550–559.

He F, Gao C H, Guo G Y, et al. 2019. Maize annexin genes ZmANN33 and ZmANN35 encode proteins that function in cell membrane recovery during seed germination. Journal of Experimental Botany. 70, 1183–1195.

Huh S M, Noh E K, Kim H G, et al. 2010. Arabidopsis Annexins AnnAt1 and AnnAt4 Interact with Each Other and Regulate Drought and Salt Stress Responses. Plant and Cell Physiology. 51, 1499–1514.

Ijaz R, Ejaz J, Gao S H, et al. 2017. Overexpression of annexin gene AnnSp2, enhances drought and salt tolerance through modulation of ABA synthesis and scavenging ROS in tomato. Scientific reports. 7, 12087.

Jain M. Khurana J P. 2009. Transcript profiling reveals diverse roles of auxin-responsive genes during reproductive development and abiotic stress in rice. FEBS Journal. 276, 3148–3162.

Jami S K, Clark G B, Ayele B T, et al. 2012. Identification and characterization of annexin gene family in rice. Plant Cell Reports. 31, 813–825.

Kim D, Paggi J M, Park C, et al. 2019. Graph-based genome alignment and genotyping with HISAT2 and HISAT-genotype. Nature Biotechnology. 37, 907–915.

Konopka-Postupolska D, Clark G, Goch G, et al. 2009. The Role of Annexin 1 in Drought Stress in Arabidopsis. Plant Physiology. 150, 1394–1410.

Kovács I, Ayaydin F, Oberschall A, et al. 1998. Immunolocalization of a novel annexin-like protein encoded by a stress and abscisic acid responsive gene in alfalfa. The Plant Journal. 15, 185–197.

Laohavisit A, Brown A T, Cicuta P, et al. 2010. Annexins: Components of the Calcium and Reactive Oxygen Signaling Network. Plant Physiology. 152, 1824–1829.

Lescot M, Déhais P, Thijs G, et al. 2002. PlantCARE, a database of plant cis-acting regulatory elements and a portal to tools for in silico analysis of promoter sequences. Nucleic Acids Research. 30, 325–327.

Li X F, Zhang Q, Yang X, et al. 2019. OsANN3, a calcium-dependent lipid binding annexin is a positive regulator of ABA-dependent stress tolerance in rice. Plant Science. 284, 212–220.

Liscum E. Reed J W. 2002. Genetics of Aux/IAA and ARF action in plant growth and development. Plant Molecular Biology. 49, 387–400.

Liu J J. Ekramoddoullah A K M. 2006. The family 10 of plant pathogenesis-related proteins: Their structure, regulation, and function in response to biotic and abiotic stresses. Physiological and Molecular Plant Pathology. 68, 3–13.

Liu Y N, Yu R J, Gao Y L, et al. 2022. Expression pattern and biological functions of an annexin encoding gene MtANN2 in Medicago truncatula under salt stress (in Chinese with English abstract). Acta Prataculturae Sinica. 31, 124–134.

Livak K J. Schmittgen T D. 2001. Analysis of relative gene expression data using real-time quantitative PCR and the 2(-Delta Delta C(T)) Method. Methods. 25, 402–408.

Lu Y G, Ou Y B, Zhang J H, et al. 2012. Genomic organization, phylogenetic comparison and expression profiles of annexin gene family in tomato (Solanum lycopersicum). Gene. 499, 14–24.

Pertea M, Pertea G M, Antonescu C M, et al. 2015. StringTie enables improved reconstruction of a transcriptome from RNA-seq reads. Nature Biotechnology. 33, 290–295.

Prochazkova D, Sairam R K, Srivastava G C, et al. 2001. Oxidative stress and antioxidant activity as the basis of senescence in maize leaves. Plant Science. 161, 765–771.

Qiao B, Zhang Q, Liu D L, et al. 2015. A calcium-binding protein, rice annexin OsANN1, enhances heat stress tolerance by modulating the production of H O. Journal of Experimental Botany. 66, 5853–5866.

Rescher U. Gerke V. 2004. Annexins - unique membrane binding proteins with diverse functions. Journal of Cell Science. 117, 2631–2639.

Shani E, Salehin M, Zhang Y Q, et al. 2017. Plant Stress Tolerance Requires Auxin-Sensitive Aux/IAA Transcriptional Repressors. Current Biology. 27, 437–444.

Singh A, Jha S K, Bagri J, et al. 2015. ABA inducible rice protein phosphatase 2C confers ABA insensitivity and abiotic stress tolerance in Arabidopsis. PLoS One. 10, e0125168.

Wang Y, Tang H, Debarry J D, et al. 2012. MCScanX: a toolkit for detection and evolutionary analysis of gene synteny and collinearity. Nucleic Acids Research. 40, e49.

Wei H, Chen J X, Zhang X Y, et al. 2024. Comprehensive analysis of annexin gene family and its expression in response to branching architecture and salt stress in crape myrtle. BMC plant biology. 24, 78.

Wei H, Movahedi A L, Liu G Y, et al. 2022. Genome-Wide Characterization and Abiotic Stresses Expression Analysis of Annexin Family Genes in Poplar. International Journal of Molecular Sciences. 23, 515.

Wu X X, Ren Y, Jiang H L, et al. 2021. Genome-Wide Identification and Transcriptional Expression Analysis of Annexin Genes in Capsicum annuum and Characterization of CaAnn9 in Salt Tolerance. International Journal of Molecular Sciences. 22, 8667.

Xie C, Zhu M, Shi R, et al. 2025. Annexins: central regulators of plant growth and stress signaling. Acta Biochim Biophys Sin. 57, 507–520.

Xu L, Tang Y, Gao S, et al. 2016. Comprehensive analyses of the annexin gene family in wheat. BMC Genomics. 17, 415.

Zhang D Y, Li J N, Niu X, et al. 2021. GhANN1 modulates the salinity tolerance by regulating ABA biosynthesis, ion homeostasis and phenylpropanoid pathway in cotton. Environmental and Experimental Botany. 185, 104427.

Zhang F, Li S F, Yang S M, et al. 2015. Overexpression of a cotton annexin gene, GhAnn1, enhances drought and salt stress tolerance in transgenic cotton. Plant Molecular Biology. 87, 47–67.

Zhao Y Z, He Z Q, Fan L J, et al. 2026. Genome-wide identification of the C2H2 zinc finger gene family in Populus euphratica and the functional analysis of PeZFP38 under salt stress. Frontiers in plant science. 17, 1754976.

